# Analysis Pipeline to Quantify Uterine Gland Structural Variations

**DOI:** 10.1101/2024.03.24.586502

**Authors:** Sameed Khan, Adam Alessio, Ripla Arora

## Abstract

Technical advances in whole tissue imaging and clearing have allowed 3D reconstruction of exocrine uterine glands deep seated in the endometrium. However, there are limited gland structure analysis platforms to analyze these imaging data sets. Here we present a pipeline for segmenting and analyzing uterine gland shape. Using this segmentation methodology, we derive individual metrics to describe gland length, shape, and branching patterns. These metrics are applied to quantify gland behavior with respect to organization around the embryo and proximity of each individual unit to the uterine lumen. Using this image analysis pipeline we evaluate uterine glands at the peri-implantation time points of a mouse pregnancy. Our analysis reveals that upon embryo entry into the uterus glands show changes in length, tortuosity, and proximity to the uterine lumen while gland branch number stays the same. These shape changes aid in reorganization of the glands around the site of embryo implantation. We further apply our analysis pipeline to human and guinea pig uterine glands, extending feasibility to other mammalian species. This work serves as a resource for researchers to extract quantitative, reproducible morphological features from three-dimensional uterine gland images in order to reveal insights about functional and structural patterns.

## INTRODUCTION

Three-dimensional (3D) imaging has paved the way for detailed study of shape and structure heretofore unavailable from conventional two-dimensional histology. Imaging technologies such as light-sheet fluorescence and confocal microscopy have yielded large, high-resolution 3D datasets containing novel findings about structure and function, particularly for branched epithelial organs including mammary glands, lungs, kidneys, and prostate glands (Chan et al., 2017; Isaacson et al., 2020; Lloyd-Lewis, 2020; Ochoa et al., 2018; Puelles et al., 2021). Such 3D reconstruction and quantitative structure analysis has allowed for the precise study of changing shape, effects of hormones and genetic perturbations on the structure and function of the dynamically developing organ.

Quantitative analysis has thus far been performed for branching in the adult mouse lung (Hwang et al., 2013), embryonic kidney (Short et al., 2010), and the mammary gland (Blacher et al., 2016; Stanko et al., 2015) and for coiling in the cochlea (Iyer et al., 2018). Quantitative study of the lung structure has revealed that branching asymmetry in the terminal bronchioles affects air mixing and subsequent blood oxygen saturation. Studies that linked lung structure to pulmonary function relied on geometric parameters such as airway diameter, length, and branching angle (Van de Moortele et al., 2018). Short et. al developed a TreeSurveyor tree segmentation method to analyze branching morphogenesis in the developing kidney and found that *Tgfb2* haplo-insufficiency increases gland bifurcation angles and reduces leading edge tip diameter (Short et al., 2013). Stanko et al, quantified branching density of mammary glands using Sholl analysis, a neurobiology method for quantifying the arborization of dendrites, and found that branching density is reduced when rats are treated with ethinyl estradiol (Stanko et al., 2015).

The International Mouse Phenotyping Consortium has used 3D quantitative imaging to phenotype 20,000 single-gene knockout mice, establish anatomical baselines, and outline statistically significant structural abnormalities associated with specific gene knockouts (Wong et al., 2012; Wong et al., 2014). In particular, as an application in branched organs, this analysis revealed that *Tcf21* mutants had significantly smaller kidneys and *Satb2* mutants had significantly smaller submandibular glands due to reduced branching morphogenesis.

3D imaging has been successfully applied to the mammalian uterus to capture various aspects of uterine glands. Recent discoveries include the developmental trajectory of mouse uterine glands (Vue et al., 2018), glandular reorganization around the embryo attachment site in the mouse uterus (Arora et al., 2016) and the existence of a plexus glandular network in the human endometrial basalis (Yamaguchi et al., 2021). In context of function and pathology, Yamaguchi et al, qualitatively discuss the appearance of human endometrial glands in patients with adenomyosis as an “ant colony-like network.” They go on to describe a “less remarkable” endometrial structure in one patient undergoing GnRH agonist treatment (Yamaguchi et al., 2021). While imaging and some quantitative analysis has been performed, uterine gland biology would benefit from identification and quantitation of novel morphological features associated with physiologic and pathologic changes, especially those that originate during the thus far elusive window of embryo implantation.

Many machine learning and artificial intelligence methods, now used to great effect in many areas of biomedical research, require quantitative metrics (“features”) as input data. Quantitative, reproducible predictions lend objective structure to subjective diagnostics that may suffer from low inter-observer reliability and poor performance. Applications of tissue-level quantitative metrics such as shape, nucleus size, epithelial volume, density, tissue grayscale intensity, and tissue texture extracted from histology and radiology, have demonstrated promise for classification of disease and prognosis in prostate (Chatterjee et al., 2015; McGarry et al., 2018), lung (Alvarez-Jimenez et al., 2020), rectal (Shao et al., 2020), and brain cancers (Rathore et al., 2019). Beyond diagnostics, unsupervised methods can be used to discover underlying patterns from unstructured data which inform descriptive models of biological systems (Gilpin et al., 2020). For example, an unsupervised model learned parameters from neural recordings that could then predict the hand velocities of a macaque during an arm-reaching task. Patterns within the raw neuron firing data were extracted and used to correctly infer aspects of the organism’s behavior (Pandarinath et al., 2018). The first step in leveraging the power of machine learning approaches is developing methods to extract features and quantitative metrics from image data sets.

Metrics that summarize structure and shape have been useful in numerous biological domains. For example, dendrite branching has long been known as a marker for neurologic disorders, such as autism (Raymond et al., 1995), Rett syndrome (Armstrong et al., 1998), Alzheimer’s (Hanks and Flood, 1991), Down syndrome (Becker et al., 1991), and chronic stress (Radley et al., 2004). Branching density, angle, and segment length have been used to characterize mammary gland (Stanko and Fenton, 2017), lung (Yu et al., 2019), and kidney morphogenesis (Sampogna et al., 2015). Tortuosity exists as a structural adaptation in organs that require higher surface area for fluid reabsorption and secretion in a limited space, such as sweat glands (Sato, 1973) and kidney (Au - McCampbell et al., 2014). Meibomian gland tortuosity, specifically, has been indexed to functional parameters such as eye dryness (tear breakup time) and has been proven to be effective as a diagnostic marker for obstructive meibomian gland dysfunction (Lin et al., 2020).

For extracting metrics from branched organs previously developed algorithms include ImageJ Simple Neurite Tracer (Longair et al., 2011), Vascular Modelling Toolkit (Antiga et al., 2008), and TreeSurveyor (Short et al., 2013). These methods are well suited for structural studies of a single “tree” that typically includes a single trunk with a number of connected complex branched structures (example mammary gland). That said, methods that target the use case of a “forest”, that includes a large population of simple branched structures (uterine glands) are limited.

In this work, we use a combination of image analysis software Imaris (Bitplane, Oxford Instruments, Abingdon UK) and Image J (Schneider et al., 2012) along with custom MATLAB (Mathworks, Natick, MA) scripts to develop an image analysis pipeline for the extraction of quantitative ‘features’ for uterine glands. We provide guidance on pre-processing tools to enhance images, model gland structure using manual and automated pipelines and extract morphological metrics pertaining to shape and branching. Where possible, to avoid spatial bias, we obtain quantitative metrics throughout the uterine horn. Our data suggests that uterine glands remodel during early pregnancy. As the embryo approaches implantation branching plateaus but glands display appreciable changes in length, tortuosity, proximity to the uterine lumen and the site of implantation. We demonstrate proof of concept of these metrics through application to murine uterine glands at different times in pregnancy, and in a single sample of non-pregnant guinea pig and human uterine glands.

## METHODS

### Animals

All animal research was carried out according to the guidelines of the Michigan State University Institutional Animal Care and Use Committee. 7-10 weeks old CD1 mice were sourced from Charles River. Pregnant mice were dissected at: GD2.5 (pre-embryo entry), GD3.5 (post-embryo entry/pre-attachment), GD4.0 (post-attachment) and GD4.5 (implantation), where the day of vaginal plug is identified as GD0.5. The guinea pig female used in this study was a 5-week-old Hartley albino strain and was sexually mature and obtained from Dr. Jason Bazil at Michigan State University. We euthanized the guinea pig using a guillotine while deep under anesthesia achieved with 5% isoflurane. Once euthanized, uteri were dissected and fixed in DMSO:Methanol (1:4) and stored at −20 degrees C.

### Human Samples

Human endometrial samples were obtained from the NIH UCSF Human Endometrial Tissue and DNA Bank. These samples were collected from women undergoing hysterectomy for non-malignant indications. Samples were collected under approved Institutional Review Board protocols. Informed consent was obtained from all patients prior to study. Endometrial samples were confirmed to be from the stratum functionalis based on visual comparison to images presented by (Yamaguchi et al., 2021).

### Whole-Mount Immunofluorescent Staining

Whole-mount immunofluorescent imaging was performed as described in (Arora et al., 2016). Briefly, uteri were dissected and fixed in DMSO:Methanol at a 1:4 ratio and stored at −20°C. For staining, samples were rehydrated in a 1:1 solution of Methanol:PBST (PBS+1%Triton) for 30 minutes followed by a wash in PBST alone for 30 minutes. Samples were then blocked in 2% powdered milk in PBST for two hours at room temperature. Uteri were then incubated with primary antibodies diluted 1:500 in blocking solution and incubated for five nights at 4°C. Uteri were then washed six times, with PBST for 30 minutes each and incubated in secondary antibodies diluted 1:500 in PBST for two or three nights at 4°C. Uteri were then washed with PBST for 30 minutes each followed by a 30-minute dehydration in 100% methanol, overnight incubation in a bleach solution of 3% hydrogen peroxide solution prepared in methanol and a final dehydration step for 30 minutes in 100% methanol. Samples were cleared using 1:2 mixture of benzyl alcohol: benzyl benzoate (BABB, Sigma-Aldrich, 108006, B6630). Primary antibodies used were CDH1 (E-CADHERIN) (M106, Takara Biosciences) for glandular and luminal epithelial staining, FOXA2 (Abcam, ab108422) for gland epithelium staining, and Hoechst (Sigma Aldrich, B2261) to stain all nuclei. The secondary antibodies used were conjugated Alexa Flour IgGs, obtained from Invitrogen Alexa-555 Donkey anti-Rabbit (A31572) and Alexa-647 Goat anti-Rat (A21247).

### Confocal Imaging

A Leica TCS SP8 X Confocal Laser Scanning microscope with white-light laser is used to image the uteri. Depending on the pipeline mode used, the objective and the interval of Z slice differs, as detailed below:

#### Global Imaging, Low Detail (GILD)

Global Imaging, Low Detail describes our imaging method for time points where we collect information across the entire uterine horn, which necessitates a lower magnification. For GILD time points, we image using a 10x objective and Z-stacks are collected at a 7.0 µm increment.

#### Local Imaging, High Detail (LIHD)

Local Imaging, High Detail describes our imaging method where we collect a high-fidelity fine structure of singular uterine glands with respect to branching dynamics. For LIHD time points, we image middle segment of the uterine horn using a 20x Objective (Leica) whose numerical aperture (NA) is similar to the refractive index of BABB and Z-stacks are collected at a 5.0 µm increment.

### Overview of image analysis pipeline

The objective of the image segmentation pipeline is to construct a centerline approximation of the uterine gland structure, enabling analysis of morphological features such as branching, tortuosity, and length.

### Image Segmentation

3D volumes of uterine glands are processed in Imaris, via Gaussian filtering operation with a sigma value of 1.44 to reduce noise (**Figure 1A**). Depending on the severity of the noise, optimally oriented flux (OOF) or Vesselness3D (V3D) filter is also applied to highlight the glandular structure relative to background. We use the Imaris Surfaces module to threshold the gland volumes, rendering them as 3D surfaces before computing the distance transform. The distance transform function converts distance from the center to intensity such that the central region of the gland where the gland lumen is present will be farthest from the edge and brightest in intensity. The transformed volumes are then used as the basis for centerline segmentation using either the A) Manual Filament, B) Automatic Filament, or C) Manual Trace pipelines.

**Figure 1:**
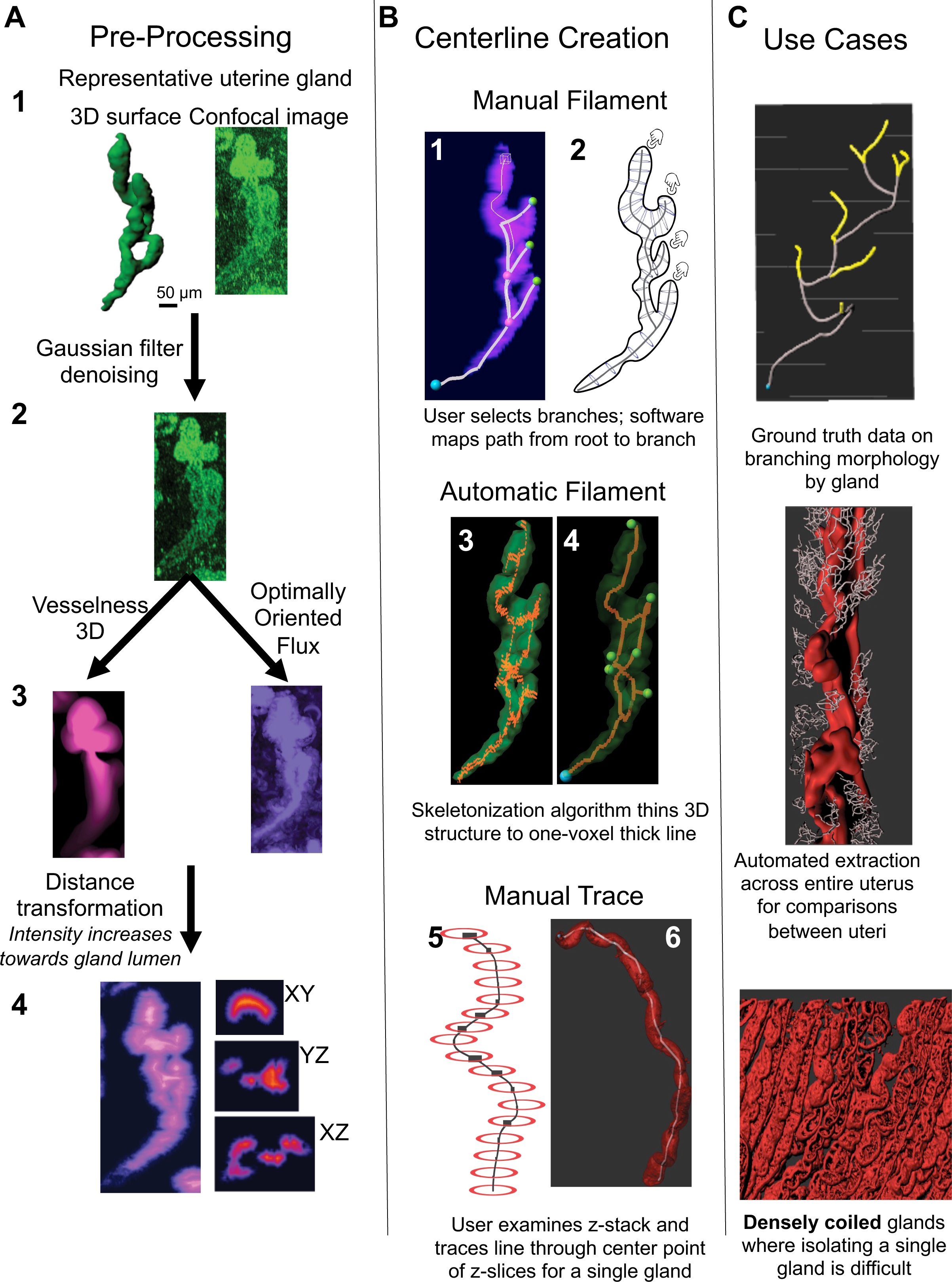
Gland Feature Extraction Pipeline. **(A)** Pre-Processing. 3D volumetric gland images are processed to remove noise using a Gaussian filter (A1). To highlight tubularity (A2) either Optimally Oriented Flux is used that enhances tubes in a low noise context, maintains branches and is prone to less error. Alternatively Vesselness3D can enhance tubes in a high-noise context. This filter tends to merge branches and is prone to more error. Distance transformation is then applied to transform the central regions of the object (gland lumen) to have the highest intensity (A3) for later centerline generation. High intensity towards the center can be confirmed using XY, YZ and ZX slicers of the distance transformed gland (A4). **(B)** Centerline Generation. Users can analyze structure through generating centerlines by Manual Filament (B1, B2), Automatic Filament (B3, B4), or Manual Trace (B5, B6). Manual Filament and Manual Trace require greatest user input but are most accurate. **(C)** Use cases for each kind of centerline generation.

### A. Manual Filament

Using the Imaris Filaments module, we semiautomatically outline branches by designating the beginning point, or root, of the gland (the point at which the gland exits the uterine lumen in 3D) and then designate the tips of the gland branches (the furthest end points of the gland away from the luminal epithelium) (**Figure 1 B1, B2**). The software calculates and constructs the shortest distance between the root and the tip while passing through the regions of highest intensity (which are the center due to the Distance Transform step). For LIHD time points, we image and select glands in a small region of the uterus. For GILD time points, we select glands regularly spaced across the entire uterine horn for an even spatial sampling.

### B. Automatic Filament

Automatic Filament describes the use of Imaris Filaments’ module with the Threshold (loops) method under automatic creation. The user can access this by creating a new Filament, then selecting “Threshold (loops)” from the “Algorithm Settings” dropdown. The distance transformed 3D volumes are passed through the Skeleton3D MATLAB script (Kollmannsberger et al., 2017) to construct one-pixel thick centerline skeletons (**Figure 1 B3**). These skeletons are then re-imported into Imaris. The threshold mode from the Imaris Filaments module is then applied to the skeletons to automatically generate centerline segmentations that contain structural data for the glands (**Figure 1 B4**).

### C. Manual Trace

Manual Trace refers to the use of Imaris Filaments’ module “Manual” method for non-automatic creation of Filaments on glands that are difficult to separate from one another. The user can access this by opting to “skip automatic creation, edit manually” and select the “Manual” method among the options listed (“AutoPath”, “AutoDepth”, “Manual”). We use Manual Trace to trace short segments connecting the centers of z-slices evenly spaced across an entire gland (**Figure 1 B5**).

### Extraction of the branch type metric

At the conclusion of each of the Manual Filament, Automatic Filament, and Manual Trace steps, we acquire a centerline segmentation in Imaris. This segmentation is represented internally in Imaris software as a series of points connected by line segments (a graph). We use a custom MATLAB script to transform the graph data structure into a tree, where nodes (branch or terminal points) in the graph contain information about the direction of the gland from root to tip (**Supplementary Code 1**). Using this new data structure, we can identify the different branch types (structural, side or terminal branch).

For each time point, 150 glands are collected from each of 3 mice, for a total of 450 glands per time point. For the prominence and reorient metrics, we use Automatic Filament to segment all the glands present in the horn.

Please refer to supplementary methods for additional preprocessing steps.

### Metrics Extracted from Image Analysis Pipeline

Centerline approximations generated using Imaris Filaments module yield a number of structural features (**Figure 2A**). In general fidelity increases with high resolution imaging and methods with user input (manual methods). Lower resolution imaging and more automated methods allow sampling of a much larger region and are efficient but cause loss in accuracy. Thus, the choice of segmentation method is determined by the set of features that are desired (**Figure 2B, 2C**). Of the listed metrics, Path Length, Branch Lengths, Branch Number, Tortuosity, and Span can be directly derived from Imaris Filaments statistics as described below. Custom MATLAB scripts were used to calculate Straight Length, Branch Types, Prominence, and Reorient.

**Figure 2:**
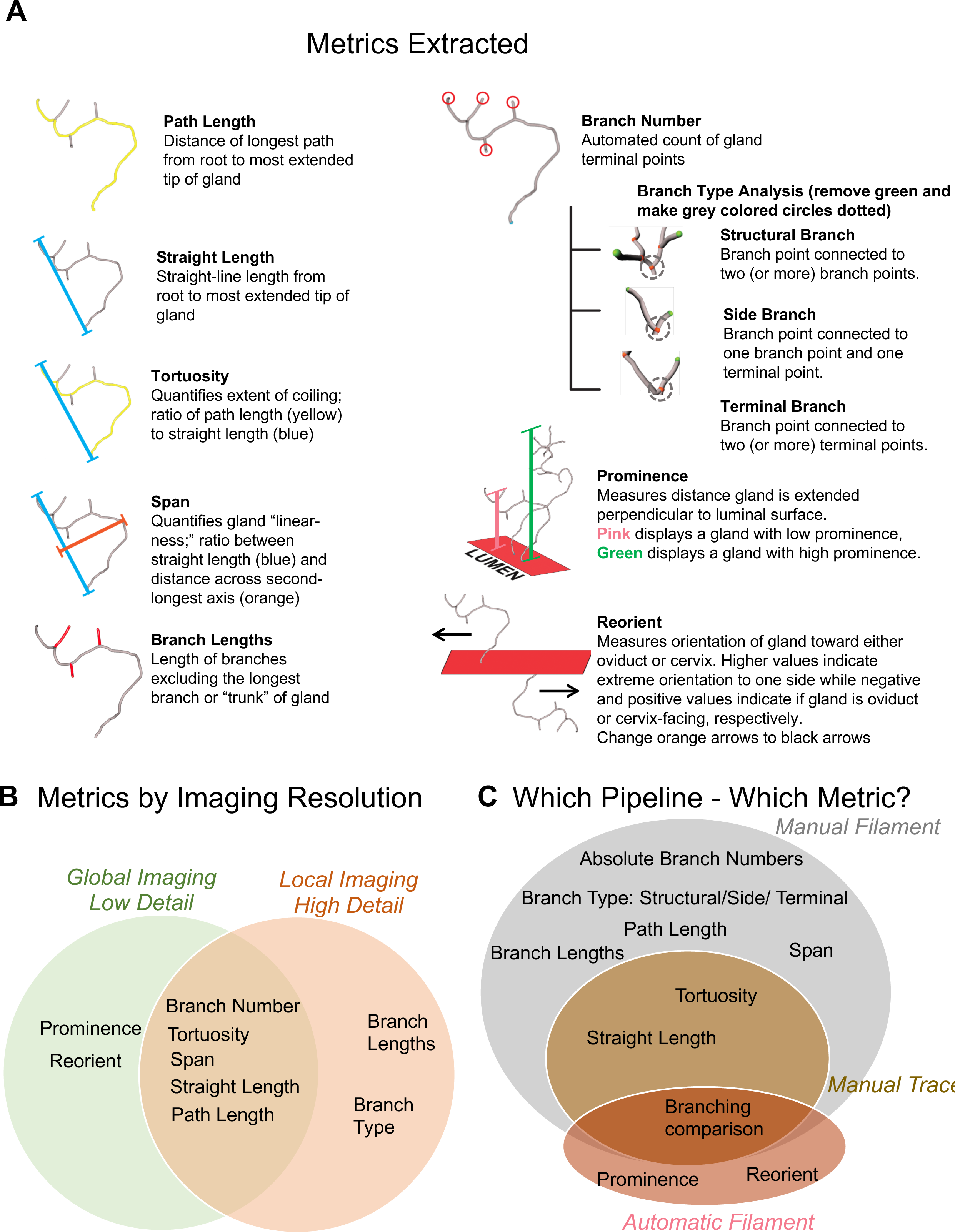
Metrics that Describe Gland Morphology. **(A)** Graphical description of different features extracted. All features except prominence and reorient are descriptive of the fine structure of individual glands, while prominence and reorient are useful for describing gland behavior across a wider set of glands throughout the uterine horn. **(B)** Local Imaging, High Detail technique is able to capture detailed branching type metrics accurately while Global Imaging, Low Detail technique can measure capture prominence and reorient metrics as it captures all glands across the entire horn. **(C)** Manual Filament and Manual Trace are gold standard methods that are able to capture any individual gland metric accurately due to the active involvement of the human user. Automatic Filament allows the capture of a large amount of data (many glands) in a relatively short period of time, but contains some error due to generation of false branches, or hyperbranching during the skeletonization process.

Metrics are defined below:

- Branch number: The number of branches is a metric extracted from the Imaris Filaments output and is reported as “Filament No. Dendrite Terminal Pts” (Bitplane, 2018).
- Branch Types: In our study we identified three types of branching:

- Structural Branching: Structural branching represents a branch point connected to two or more branch points, with no free end points in any of the branches emerging from the branch points.
- Side Branching: In the side branching case, at least one of the branches extending from the branch point does *not* have an endpoint, indicating that the tree can continue to grow further.
- Terminal Branching: This is a case where a branch point branches into two or more branches with end points. No more branches can be generated from a terminal branch.

We extract branch types using a suite of custom MATLAB scripts (**Supplementary Code 1**) that extends Imaris Filaments functionality by translating the Filament to a Tree data structure (Black, 2017) A Tree is comprised of Nodes, which contain Imaris statistics, while the edges between Nodes capture the hierarchical relationship from the root Node (beginning point) to the terminal Nodes (branch endpoints) with branch Nodes in between (Nodes connecting Nodes). Representing a Filament as a Tree facilitates analysis of branching type, by analyzing the relationship between Nodes.

- Path Length: Path length describes the length from the gland root to the most extended tip. This can be also considered as the “trunk length,” essentially the sum of the segments beginning from the gland root to the furthest tip. This measure excludes branch lengths and captures the length of the central path through the gland. In Imaris, this metric is the maximum “Point Distance” of each Filament object (**Supplementary Code 2**).
- Straight Length: Straight length is the shortest straight-line distance from the gland root to the furthest tip. In Imaris, this metric is the Euclidean distance between the coordinates of the Beginning Point and the Terminal Point (Bitplane, 2018). The terminal point is the branch tip furthest away from the root of the gland (**Supplementary Code 2**).
- Average Branch Length: Branch length denotes the length of the branches that are *not* part of the “trunk,” or the path from the gland root to the furthest tip. We calculate the “Average Branch Length” by subtracting “Path Length” from “Filament Dendrite Length (sum)”, extracted from Imaris, and divide this value by the number of branches (reported in Imaris as “Filament No. Dendrite Terminal Pts”) (Bitplane, 2018) (**Supplementary Code 2**).
- Tortuosity: Tortuosity is the ratio between the path length and straight length. The higher the tortuosity, the more coiled the gland is, as a greater length along the body of the gland is packed into a smaller linear distance from root to tip (**Supplementary Code 2**).
- Span: Gland Span can be visualized similarly as hand or wingspan. Formally, if a rectangular box were fitted around the gland, Span is the ratio between the longest and second-longest axes of that box. In terms of Imaris metrics, Span is BoundingBoxOO Length C divided by BoundingBoxOO Length B (Bitplane, 2018) (**Supplementary Code 2**).
- Prominence: Prominence measures the gland’s angle relative to the luminal epithelial surface local to the gland’s beginning point. A custom MATLAB script **(Supplementary Code 3)** calculates this value by constructing a normal vector from the local luminal epithelial surface and comparing it to the vector between the gland’s beginning point and most extended terminal point (“root-to-tip” vector). Glands that are perpendicular to the luminal surface (where the root-to-tip vector is parallel to the luminal surface’s normal vector) have a prominence value of 1. On the other hand, glands that are parallel to the luminal surface (where the root-to-tip vector is perpendicular to the luminal surface’s normal vector) have a prominence value of 0. With this in mind, a gland’s prominence value is calculated as the proportion of the root-to-tip vector’s total length that is in the direction of the luminal surface’s normal vector.
- Reorient: Reorient measure indicates the gland’s direction along the oviductal-cervical axis and its angle with the uterine lumen. Higher reorient values indicate the gland’s angle relative to the lumen is lower (i.e: the gland faces more toward a particular direction – oviduct or cervix). Calculating the reorient value requires a custom MATLAB script **(Supplementary Code 3)** operating on an image that was aligned such that the oviductal-cervical axis was parallel to the image Y-axis. To achieve this, we split the uterine horn into parts and rotated any curved or slanted sections to be aligned with the Y-axis. After rotation, the split parts were rejoined to recreate the entire horn as a single unit. The custom MATLAB script assumes the point on the gland filament closest to the luminal epithelium is the beginning point and uses the point furthest extended on the Y-axis (in either direction) as the end point of the gland. The reorient statistic is formally the difference in the y-coordinate between the beginning point and the end point of the gland. Thus, the magnitude of the reorient value indicates the extent to which a gland “leans” in one direction or the other while the sign of the reorient value indicates the direction. A negative value indicates the gland is facing the oviduct while a positive value indicates the gland is facing the cervix.

### Statistical testing

All statistical analysis is performed in R. Significance is determined at p < 0.05, and all hypothesis testing is done via Wilcoxon Rank Sum test with Benjamini-Hochberg correction to account for non-normality of data distributions.

## RESULTS

### Local Imaging in High Detail (LIHD) reveals glands lengthening after embryos enter the uterus

The middle segment of the uterine horn was imaged at high resolution (20x magnification) and analyzed for changes in gland structure. We evaluated the early pregnant mice at gestational day (GD) 2.5 and GD3.5 stages. At GD2.5 the embryos are still in the oviduct thus this stage represents gland metrics “pre-embryo entry”. At GD3.5 the embryos are in the middle of the uterine horn (Flores et al., 2020) and thus this stage represents gland metrics “post-embryo entry”. The goal with LIHD imaging is to sample a small region of the uterus to generate ground truth data. It is technically challenging to capture the entire horn due to limitations of imaging time and file size to be processed on the image analysis software.

Using LIHD we assessed gland branching and the type of branch pattern at different stages in early pregnancy. Using the metric extraction pipeline (see methods), we converted raw image data of uterine glands into gland lumen representations (“Filaments”, **Figure 3A**). We note that total branches are similar at GD2.5 and GD3.5 (**Figure 3B1, 3B2)**. Further, we find that the frequencies of each type of branch (structural, side, and terminal) at both time points stay the same (**Figure 3B3**). When examining branch type *by individual gland* (**Figure 3C**), we find that all types of branches are again consistent between GD2.5 and GD3.5.

**Figure 3:**
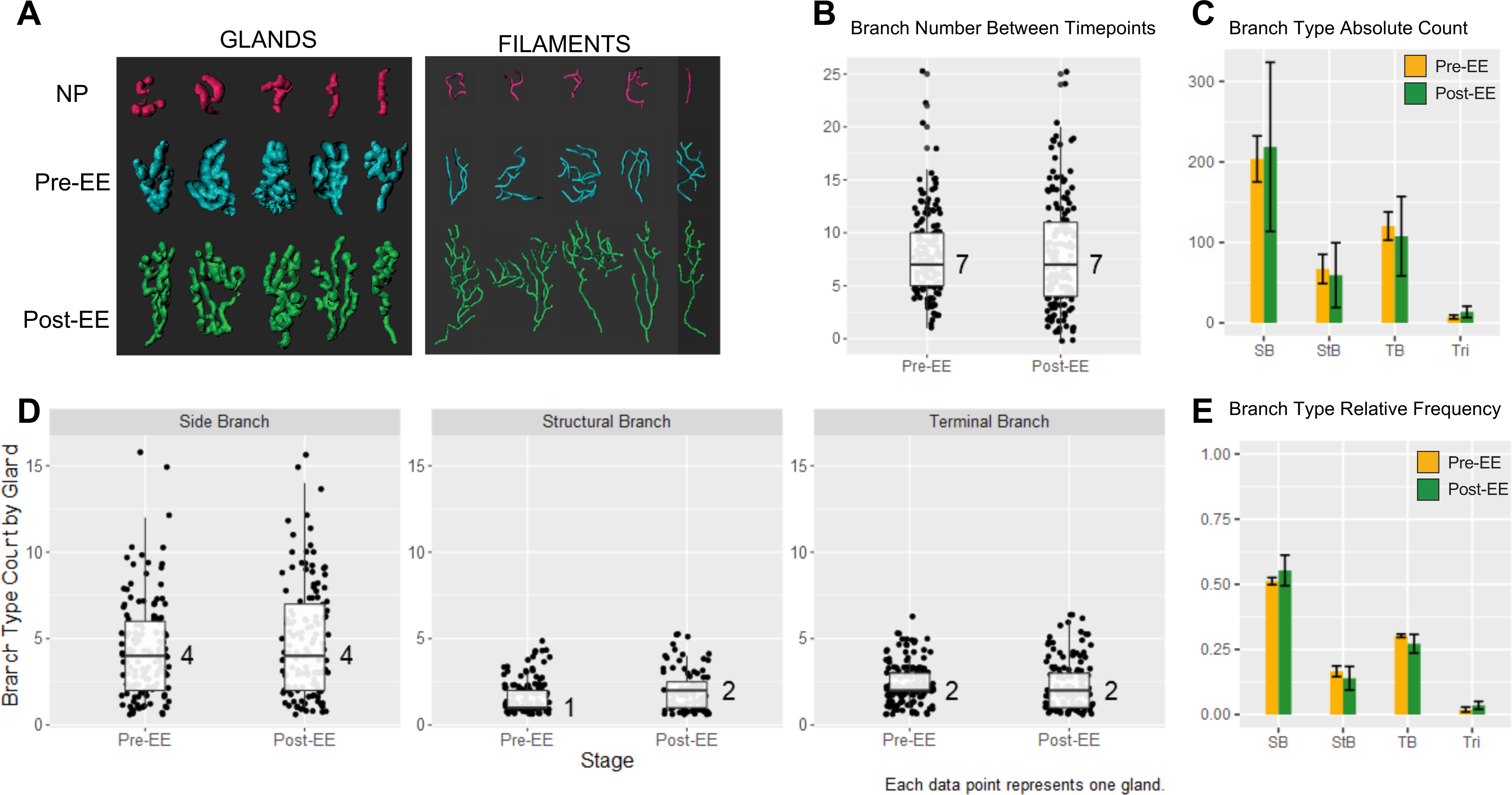
Insights from Analyzing the Branching Morphology of Uterine Glands at different stages with Local Imaging in High Detail. **(A)** Example 3D renderings that show correspondence between gland and filament structures. Changes in **(B)** raw branch number (boxplot) and **(C)** absolute number of different branching types (B2) at different stages analyzed. NP: Non-Pregnant; Pre-EE: Pre-Embryo Entry (GD2.5); Post-EE: Post-Embryo Entry (GD3.5); SB: side branch, StB: structural branch; TB: terminal branch; Tri: trifurcation. **(D)** Proportional representation of each type of branching among the sample of glands in each stage. Cases with no branches were excluded. **(E)** Relative frequency of each branch type after normalizing to total number of glands. All graphs use a total sample size of 150 glands across 3 mice with 50 glands from each mouse. All significance testing uses Wilcoxon rank-sum test.

Another aspect of uterine gland development is their linear structure. Guided by prior qualitative observations that glands lengthen during early pregnancy (Arora et al., 2016) we measure metrics for length, span and tortuosity. We note the path length (**Figure 4A**), straight length (**Figure 4B**) of glands significantly increase between GD2.5 and GD3.5. Tortuosity (**Figure 4C**) and average branch length (**Figure 4E**) is similar at GD2.5 and GD3.5. Span on the other hand decreases from GD2.5 to GD3.5 (**Figure 4C**).

**Figure 4:**
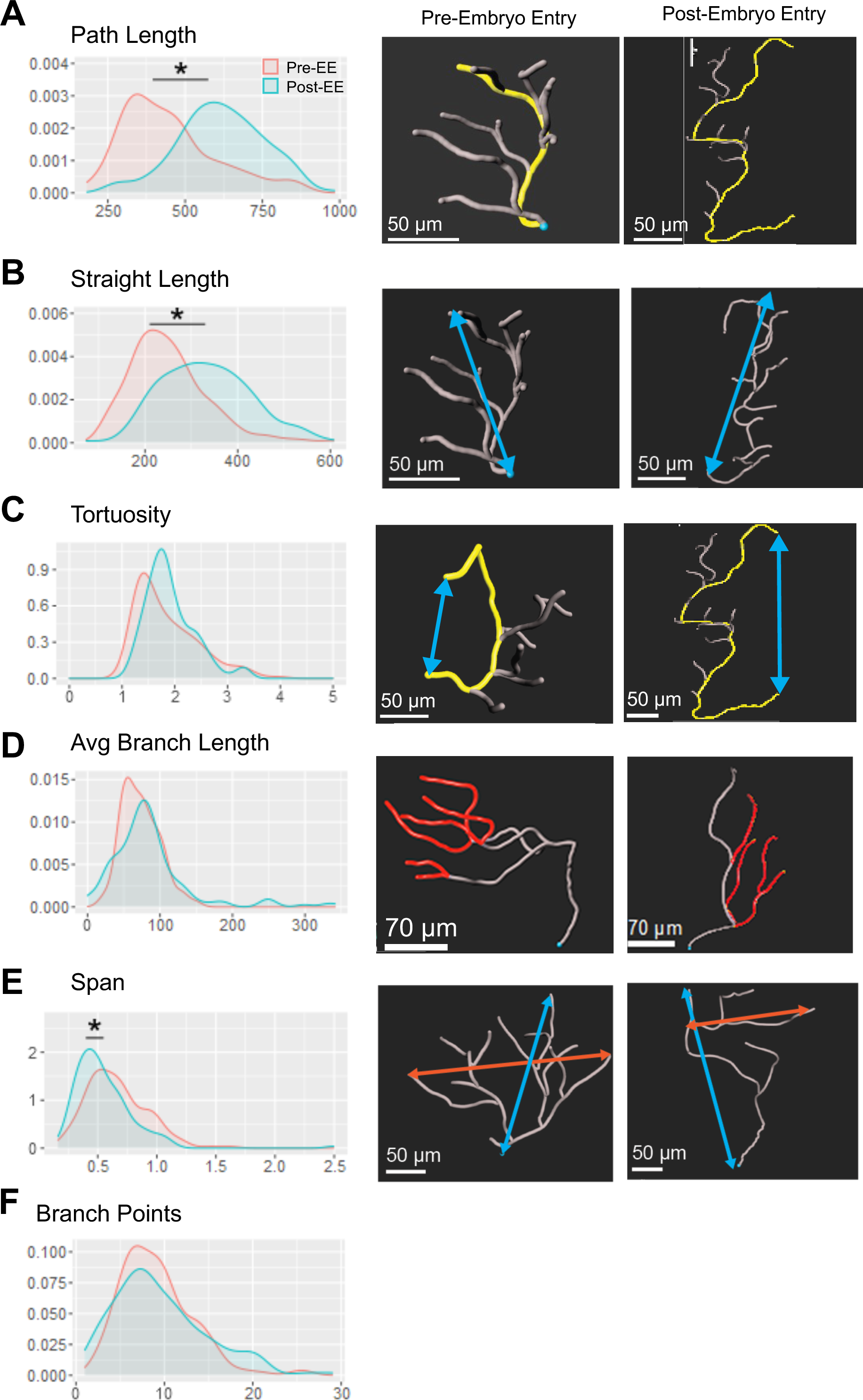
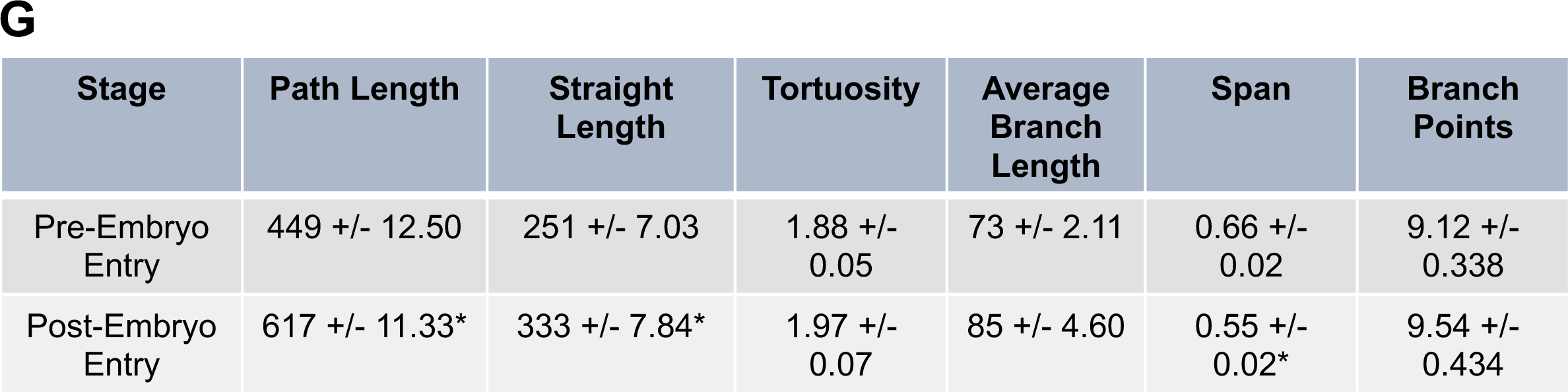
Dynamic changes in uterine gland morphology between non-pregnant uteri and early stages of pregnancy until the embryos enter the uterus. **(A)** Path length, gland’s length along the longest path from root to tip; **(B)** Straight length, a measure of the gland’s shortest length from root to tip; and **(C)** Tortuosity, a measure of coiling that is a ratio of path length to straight length all increase as embryos enter into the uterus. **(D)** Average branch length measures the lengths of branches that emerge from the gland’s “trunk,” normalized by number of branches per gland, increases from non-pregnant to early pregnant stage. **(E)** Span measures the gland’s structure in terms of extension along an axis perpendicular to the axis in the direction of the most extended tip. Increases between non-pregnant and pre-embryo entry stage but then decreases post-embryo entry. **(F)** Probability density plot comparing branch point number per gland between timepoints. **(G)** Table listing means +/- standard error of each metric across time points. Asterisk marks value as significantly different from *all other stages*. Significance testing was performed via the non-parametric Wilcoxon rank-sum test. * indicates p<0.05.

### Comparison of local imaging in high detail (LIHD) and global imaging in low detail (GILD) suggests differences in measurement of absolute metrics

While LIHD gives more detail, in order to capture metrics across the entire uterine horn we used global imaging in low detail (GILD). We then compared individual metrics using LIHD and GILD for GD3.5. We observed that while measurements for branch number, path length and span were similar at both resolutions, absolute values for straight length and tortuosity (a metric dependent on straight length) are significantly different (Supplementary Figure 1). These data suggest that to make meaningful interpretations two conditions that are being compared should be imaged at the same resolution.

### Global Imaging in Low Detail (GILD) Reveals Glands “Stretch” Toward Implantation Site prior to Embryo Implantation

We used GILD to sample glands across the entire horn. We analyzed changes in gland structure between GD3.5 (post-embryo entry but pre-embryo attachment) and GD4.0 (embryo attachment time point). During attachment, GD4.0, glands show a similar number of terminal points when compared to GD3.5 glands (**Figure 5A**). We observed that GD3.5 glands increase in path length (**Figure 5B**), straight length (**Figure 5C)** and decrease in span (**Figure 5D**) compared to GD4.0 glands, similar to the transition between GD2.5 and GD3.5 (**Figure 4A, 4B, 4C**). Tortuosity does not change significantly with embryo attachment **Figure 5E**). Coincident with gland lengthening, glands also “bend” closer to the surface of the lumen, yielding lower prominence from post-embryo entry/pre-attachment to post-attachment (**Figure 6A**, **6B**).

**Figure 5:**
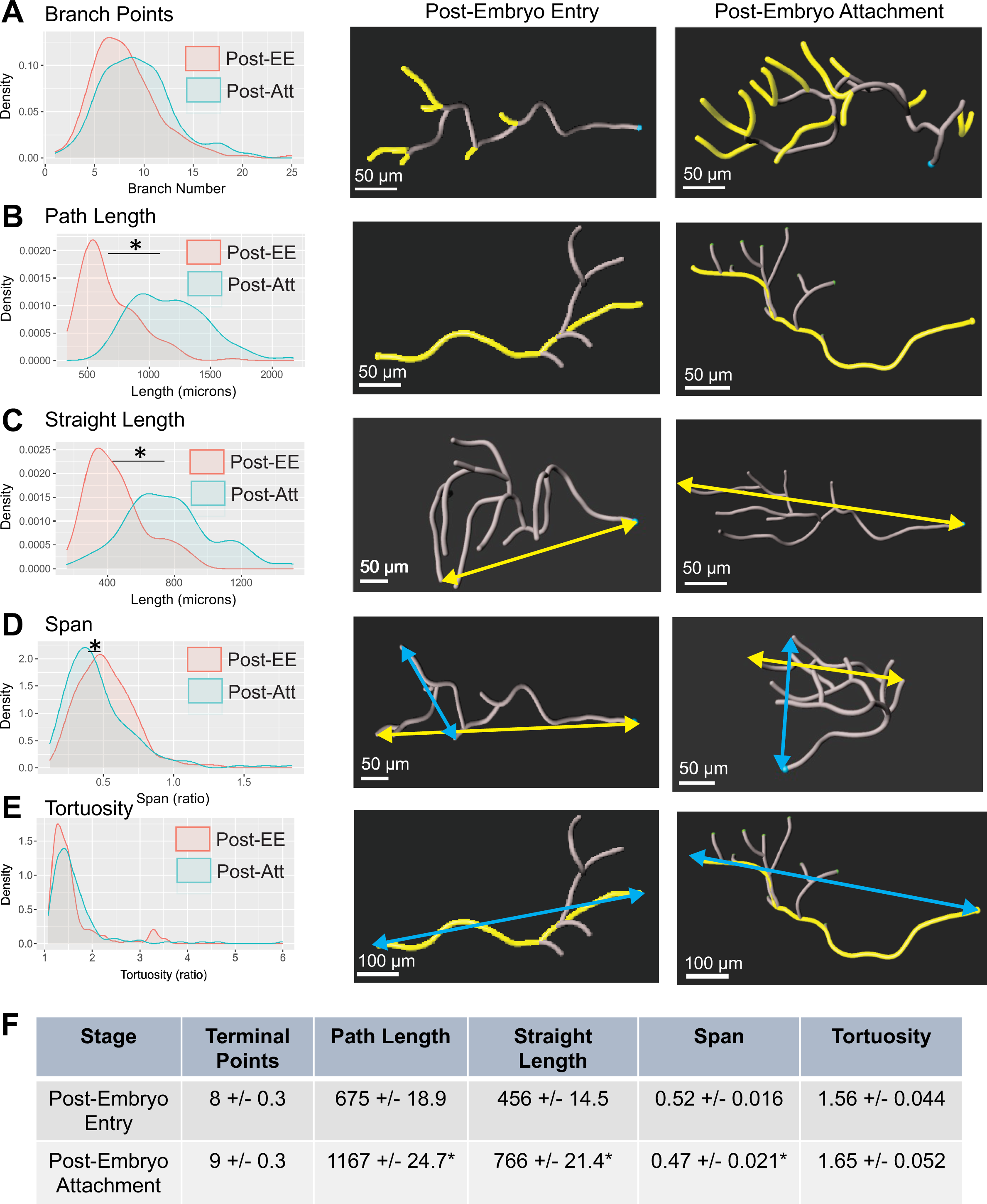
Dynamic changes in uterine gland morphology when embryos have entered the uterus and when embryos are beginning attachment. **(A)** A small increase in branching is observed. Both **(B)** Path Length and **(C)** Straight Length increase while **(D)** Tortuosity remains the same. **(E)** Span decreases closer to embryo attachment, an indication that gland branches are lengthening further “sideways.” (F) lists mean+/- standard error values of each metric across time points, asterisk marks significantly different values according to Wilcoxon rank-sum test. * indicates p<0.05.

**Figure 6:**
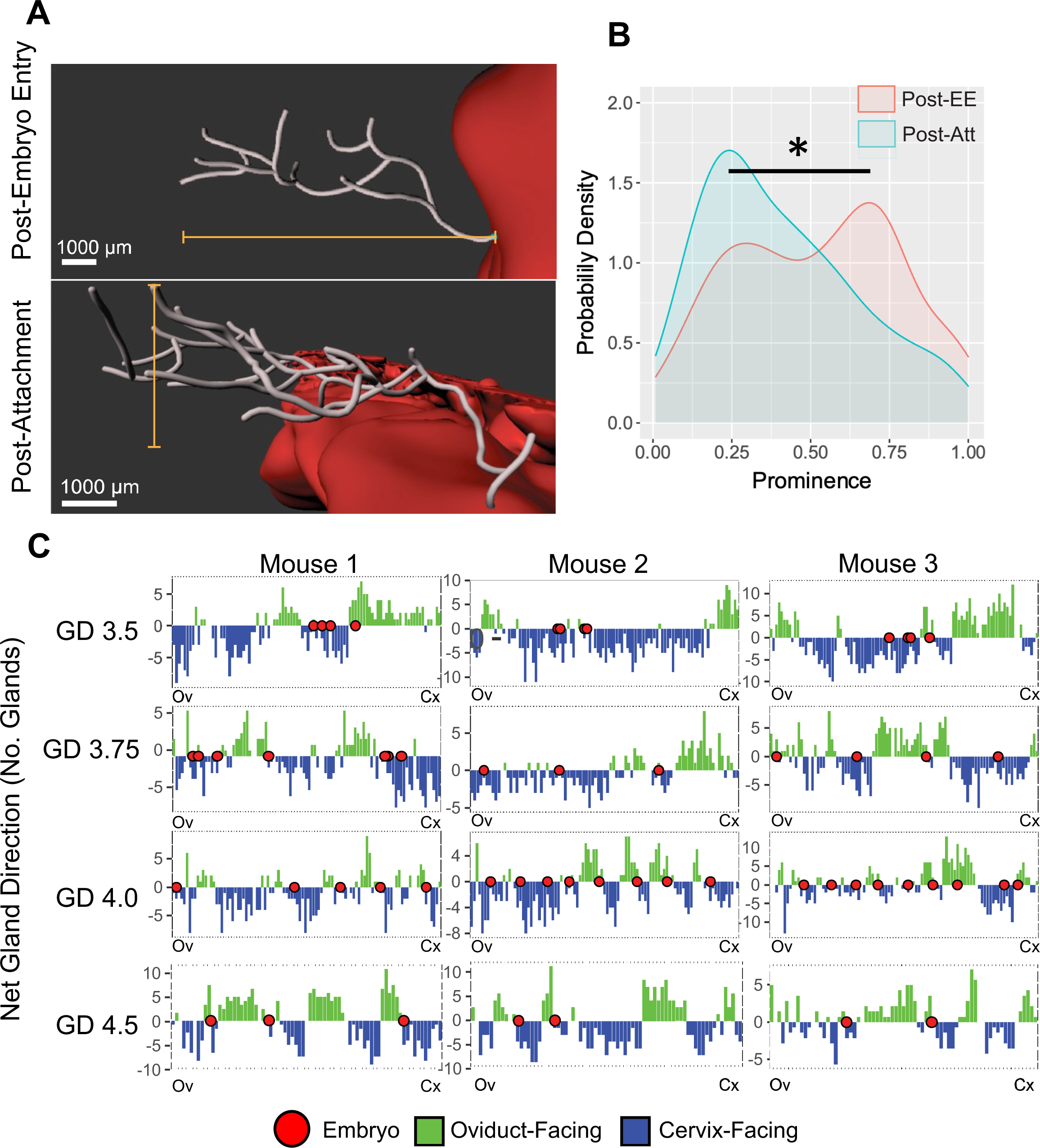
Prominence and Reorient Metrics along the entire length of the uterine horn. **(A)** Visual comparison of prominence, a measure of glands perpendicularity relative to the uterine lumen. Glands are oriented in the same direction at different stages, but the lumen is perpendicular to the gland post-embryo entry but pre-attachment (GD3.5), top compared to post-attachment (GD4.0), bottom. **(B)** Quantitative distribution of gland prominence as shown in (A). Significance testing was performed via the non-parametric Wilcoxon rank-sum test. * indicates p<0.05. **(C)** Reorient measures the direction of a gland either toward the cervix or oviduct. As embryo nears implantation, glands reorient to form sites where glands on either side of the implantation site face the oviduct and the cervix.

Glands reorient around embryo attachment sites (Arora et al., 2016). To quantify this reorientation metric, we analyze the orientation of glands along the oviductal-cervical axis (captured by the positive or negative sign of the reorient metric). Although previously reported at GD4.5, our metrics suggest that gland reorientation sites appears as early as GD3.75 with the embryos present in proximity, and by GD4.0 embryos are present in the middle of reorientation sites (Madhavan et al., 2022). By GD4.5, the time of embryo implantation, embryos are evenly spaced out at the centers of gland reorientation sites (Flores et al., 2020) (**Figure 6C**).

### Application to Other Species (Guinea Pig and Human)

To illustrate the application of our method to other species we apply our methodology to uterine glands from the guinea pig and human endometrium. Manual and Automatic Filament method can be applied to guinea pig while quantitation of the human endometrial biopsy glands requires Manual Trace due to the high density of the glands clustered tightly together, rendering it difficult to separate glands from one another digitally. Five-week-old guinea pig (**Figure 7A**) and human endometrial biopsy (**Figure 7B**) samples are analyzed using Manual Filament and Manual Trace methods, respectively. Post-pubertal guinea pig glands are comparable in structure to non-pregnant diestrus mouse glands (Supplementary Figure 2) while human glands are much longer, more coiled, and less branched (**Figure 7B**). As reported previously these glands best depict glands of the human endometrial functionalis layer (Yamaguchi et al., 2021).

**Figure 7:**
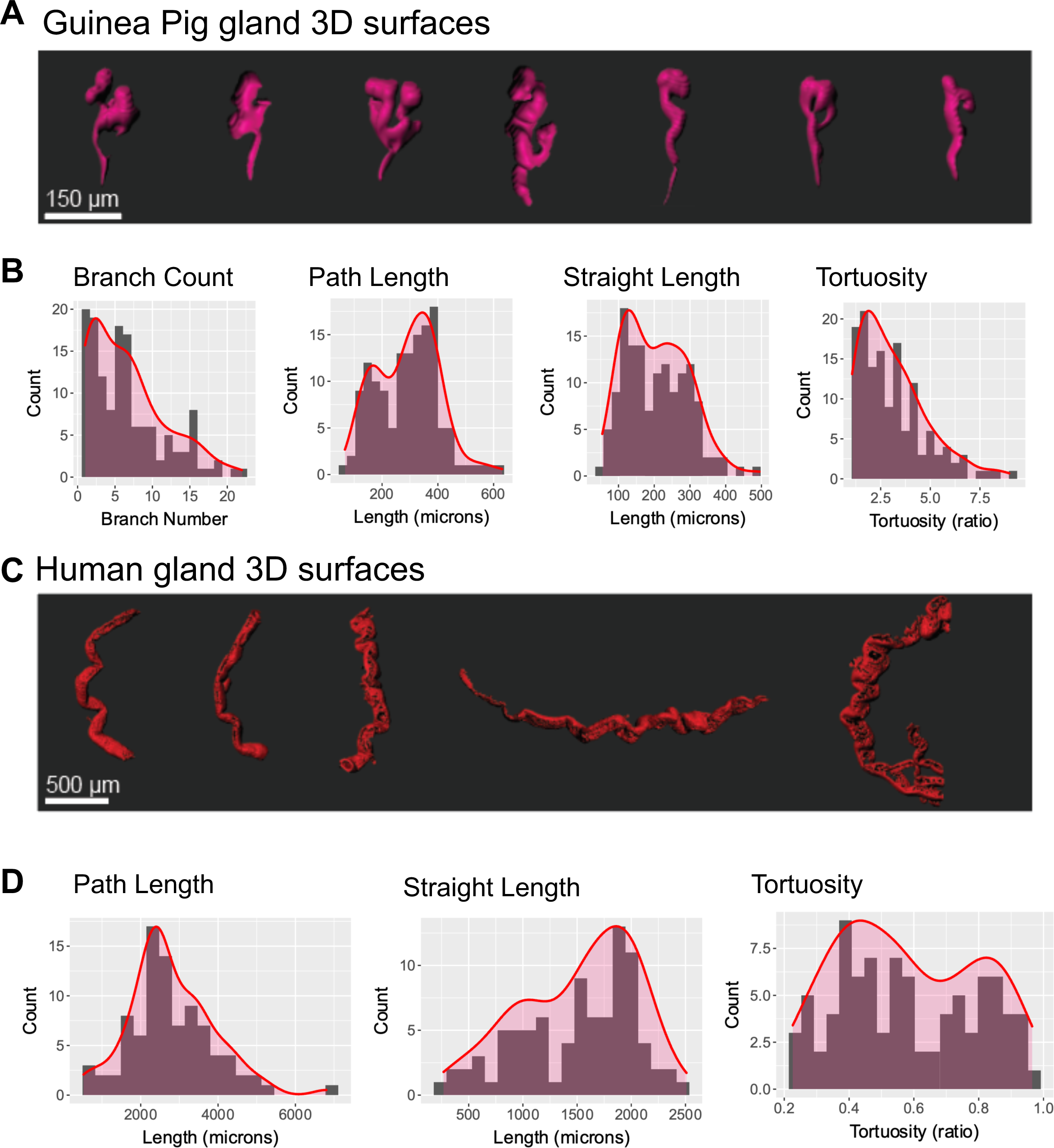
Application of gland feature extraction pipeline to Guinea Pig and Human Uterine Glands. **(A)** Gland surfaces from guinea pig imaged using Global Imaging, Low Detail. **(B)** Histogram distributions of Branch Count, Path Length, Straight Length and Tortuosity in the Guinea Pig sample. **(C)** Gland surfaces from human hysterectomy biopsy imaged using Global Imaging in Low Detail. **(D)** Histogram distributions of Path Length, Straight Length and Tortuosity for human uterine glands. Human glands are mostly unbranched thus the branching metric was excluded.

## DISCUSSION

### Technical Advances

We present an image analysis pipeline for uterine gland segmentation and analysis that can extract several biologically informative features. This pipeline is extensible to other mammalian species beyond mice, as demonstrated by descriptive results from guinea pig and human uterine glands.

### Challenge of Analyzing Forests versus Trees

Among image quantification methods for branched epithelial structures, ours is the first to address a tissue that is a “forest” rather than a “tree.” We use extracted metrics to sample, describe, and compare the behavior of the entire gland population across time points and in some cases, such as reorient, we forego sampling and are able to use metrics for every gland in the horn. Comparable analyses has been performed on a few large complex structures, such as branched and unbranched mammary glands, developing kidneys and lungs (Chan et al., 2017; Lloyd-Lewis, 2020; Ochoa et al., 2018; Puelles et al., 2021). However, unlike these branched organs that are tree structures, uterine glands are forest like. They contain much structural diversity *within* each group and between groups there is also a continuum of shape changes with appreciable overlap. Thus, differences between non-pregnant and early pregnancy stages cannot be appreciated unless a large fraction of the glands from each time point is sampled. Uterine glands vary from being highly linear, ramified structures, to being globular, cystic, and coiled. Like comparing two heterogeneous forests, comparing uterine gland populations presents the challenge of high intraclass diversity.

We address this problem in two ways:

1. We provide a methods pipeline (LIHD + Manual Filament) that generates “ground-truth” results. Branch terminal points and Filament shape are generated via user input and thus amount to manual user annotations of structure.
2. We provide evidence that lowering resolution to capture the whole horn (GILD) recapitulates some observations from ground-truth LIHD metrics. We further provide evidence that Automatic Filament offers an automated way to generate certain metrics (prominence and reorient) that match qualitative observations already established in the literature (Arora et al., 2016; Madhavan et al., 2022)

### Advantages and Limitations of Local Imaging in High Detail (LIHD)

We illustrate the utility, benefits, and caveats of several different modes of our pipeline. Local Imaging in High Detail (LIHD) allows the user to extract detailed branch information, such as branch lengths and branching type, from a spatially limited subset of glands. LIHD is best suited to study of structure at an individual level (examining a “tree”), such as illustrating branching morphogenesis. LIHD measurements further serve as a ground truth reference for validating either automated or lower-resolution methods. While LIHD measurements are highly accurate, the required imaging resolution generates very large file sizes (on the order of tens of gigabytes) for a small subsection of the uterine horn. Storing LIHD captures of the entire uterine horn would require more data storage and it would be unfeasible to process using image analysis software such as the current version of Imaris (v9.2.1), precluding any metrics describing gland structure across the entire length of the uterus.

### Advantages and Limitations of Global Imaging in Low Detail (GILD)

Global Imaging in Low Detail (GILD) enables population-level metrics that quantify characteristics of the “forest” of glands across the uterine horn, for reorientation around the implantation site and proximity to the lumen. We use GILD to capture information about glands relative to proximity to the embryo. GILD is well-suited for such use cases, which require information about glands across the entire horn, and not limited to a sample drawn from a single location. Making conclusions about the behavior or structure of *uterine glands as a group biological unit* (“forest”) requires spatially unbiased sampling across the entire horn, necessitating the use of GILD, whereas conclusions about the individual structure of glands (“tree”) can be informed by measurements from a spatially limited sample, LIHD.

For metrics that overlap between LIHD and GILD, namely branch number, tortuosity, span, straight length, and path length, the GILD pipeline would produce different results than the ground-truth LIHD pipeline if the user were to analyze the same sample of glands with both. We observe that the low resolution of GILD caused branches in close proximity to merge with each other and appear as one branch. GILD’s lower resolution also reduces tortuosity and path length metrics: this is likely because features such as coils and twists are reduced to lines when images are captured at a lower resolution. However, we show that some GILD-generated metrics at uterine embryo entry pregnancy time point are similar to LIHD-generated metrics. This consistency across both LIHD and GILD, establishes GILD-generated metrics as accurate for *relative* comparisons between time points. This relative accuracy makes a combination of GILD and LIHD metrics applicable for studies analyzing differences in stage, genetic phenotypes, inhibitor effects, or other biological conditions. Although we have demonstrated the *relative* accuracy of GILD metrics on our murine dataset, we still suggest researchers using GILD validate their measurements against a smaller LIHD dataset to ensure consistency of differences between biological conditions.

### Semi-Automatic versus Automatic Segmentation

Our pipeline describes a semi-automatic mode (“Manual Filament”) and an automatic mode (“Automatic Filament”) for segmentation. Manual Filament can capture all metrics accurately but requires extensive user input, about four hours for fifty glands. Automatic Filament requires little user input and can generate prominence and reorient metrics for the entire population of glands. Automatic Filament requires little user input, about 15 minutes of active user attention for one uterine horn. Actual time taken will vary based on available processing power and the dataset file size; Imaris Threshold Filaments requires the bulk of time and processing resources in the pipeline. The drawback of Automatic Filament is the highly inflated branch numbers. The skeletonization algorithm used in this pipeline interprets small, random protrusions on the gland surface as branches, thereby overestimating branch number. While Automatic Filament is uninformative for analyses of branch points in an individual gland, it is still viable for prominence and reorient, since these metrics only require an accurate estimation of the gland’s beginning point and furthest extended point.

### Biological Advances

Uterine glands are required for embryo nourishment during early stages of mammalian pregnancy prior to placenta formation (Kelleher et al., 2019). Thus, changes in uterine gland structure during embryo movement and implantation may inform us of uterine gland’s function. Most of the studies on uterine epithelium have thus far focused on the changes in the luminal epithelium (LE) since the embryo first makes contact with the LE. However, using 3D reconstruction and segmentation we are able to document changes in the glandular epithelium with respect to shape, branching and proximity to the LE as the embryo approaches implantation.

For uterine gland branching analysis, individual metrics by themselves will be less informative than the holistic view offered by combining multiple metrics together. Uterine gland branching does not change between uterine pre-embryo entry and uterine post embryo entry time points. Increased epithelial proliferation during early pregnancy is a result of estrogen that stimulates epithelial growth (Hewitt et al., 2003; QUARMBY and KORACH, 1984; Wang et al., 2000). Since estrogen levels rise and fall prior to embryo entry into the uterus gland branch number should remain unaffected as embryos enter the uterus and our quantitative analysis support this idea. The role of cyclic hormones in increase in the number of glandular epithelial cells was established in a previous study (Jin, 2019). Further we have recently shown that epithelial-specific deletion of estrogen receptor results in glands that fail to branch (Granger et al., 2023) suggesting epithelial estrogen signaling is a driver of gland branching.

Our data suggests that the transition between pre and post-embryo entry is profiled by an increase in path length and straight length and a decrease in span without affecting average branch length. Further, the transition between embryo entry to attachment, is profiled by an increase in path length, straight length, tortuosity, and a further decrease in span. This period coincides with the rising progesterone levels from the corpus luteum, causing cessation of epithelial proliferation (Wu et al., 2018). Our quantitative findings match qualitative reports in the literature with regard to cessation of branching (Wu et al., 2018) and suggest that progesterone stimulates gland lengthening and tortuosity in preparation for embryo attachment and implantation chamber formation. We hypothesize that these changes are driven by a progesterone dominant environment resulting in gland elongation and stretch, either due to cell autonomous changes in the glands or changes in the stromal cell signature and extracellular matrix environment that impact the gland structure. These ideas will be tested in future studies using hormone receptor knockouts. The prediction would be that progesterone receptor depletion either in the epithelium or the stroma will fail to show changes in gland shape without affecting branch numbers.

### Glands Exhibit Structural Adaptations to Optimize for Functional Stimulus

Uterine glands are considered exocrine glands: secretions are generated and expelled through the apical surface into the gland lumen to be delivered into the uterine lumen. Some studies suggest that uterine glands secrete paracrine decidualizing factors into the stroma (Filant and Spencer, 2014), implying secretions are produced basally and delivered to the nearby cells. Increase in gland lengthen and coiling toward implantation sites as they undergo gland reorientation (Arora et al., 2016) would support a paracrine mechanism of secretion. Gland reorientation coincides with formation of flat preimplantation regions in the lumen where the center eventually becomes the implantation site for an embryo (Madhavan et al., 2022). This indicates that P4-mediated stretch could be a mechanism to pull on the luminal epithelial folds to create the flat implantation region. The gland stretch could also be an adaptation to bring gland secretory sites (for basal secretion) closer to the location of the stroma surrounding the implantation chamber for decidualization. Further, prominence decreases closer to attachment suggesting that the glands are in close proximity to the luminal epithelium.

This may be a functional adaptation to bring basolateral secretion Leukemia inhibitory factor (LIF) from the uterine glands closer to basally expressed LIF receptors on the luminal epithelium to enable uterine receptivity (Cheng et al., 2017). Such uterine gland adaptations for embryo implantation are reminiscent of structural adaptations in the sweat glands where increased tubular length and volume of the glands are observed in response to hotter climates and exercise (Sato et al., 1990).

## Limitations of the study

### Staining

The quantitation method relies on good quality immunostaining. Poor staining can result from user error or tissue conditions. The lumen at the estrus stage is very fluid-filled and tissue at later gestational stages becomes thicker. Both of these conditions affect the penetrative ability of primary and secondary antibodies and thus may result in high noise, holes, and disconnected branches from the gland structure. Preprocessing filters can partly address lower-quality images; however, poor staining quality can still affect several metrics.

Prominence and reorient are affected by sparse staining since these measurements are not evenly spatially sampled and rely on the density of glands throughout the horn. If certain regions of the horn have poor staining and thus have less glands imaged in that region, prominence and reorient metrics are less informative for describing gland reorientation relative to uterine region, embryo movement or other uterine environmental processes. Branch points, branching type, and average branching length measures are less informative since sparse staining often results in disconnected branches. The Manual Filament method relies on the user to recognize that a disconnected branch may belong to a gland and thus adds uncertainty to whether gland structures being described are complete or not. Branching measure are also affected by high image noise, which exacerbates the problem of merging separate gland branches in proximity. Path length, straight length and derivative metrics (tortuosity, span) may also be affected if staining is of poor quality where glands may display incomplete or disconnected branched structure.

### Imaging Resolution

As discussed previously, due to lower resolution, GILD captures only a subset of metrics *absolutely* accurately, but retains accuracy for comparison across biological conditions. LIHD, conversely, is limited spatially but is accurate for highly detailed metrics of individual gland structure, such as branching type.

### Segmentation

This pipeline could be improved by superior denoising algorithms, specifically methods that could impute missing holes or branches in a glandular structure. The automated skeletonization script used in the Automatic Filament pipeline inflates the number of branches by registering topological irregularities on the gland surface as small branches. Even after pruning, branch numbers remain high. An improved skeletonization method that could overcome this challenge would further improve the automation potential of this pipeline.

## Conclusion

Generation of quantitative metrics combined with qualitative observations from the existing literature allowed us to propose several possible structural adaptations to enhance uterine gland function. We believe reproductive development study will benefit from a structure-function framework that uses quantitative structural data to illustrate changes in the uterine environment between 1) time points, 2) mutant phenotypes, 3) chemical treatments, or 4) artificial conditions (e.g: ovariectomy, pseudopregnancy), among other possibilities. We believe that this method represents a valuable asset for any reproductive researcher investigating how uterine environments affect uterine gland shape and function.

## CONFLICT OF INTERESTS

The authors declare no conflict of interests.

## AUTHOR CONTRIBUTIONS

S.K. and R.A. designed the experiments; S.K. performed experiments; S.K., A.A., and R.A. analyzed the data. S.K., and R.A. interpreted the results; S.K. and R.A. wrote the manuscript.

## FUNDING

We acknowledge support from March of Dimes grant #5-FY20-209.

## ACKNOWLEDGEMENTS

We acknowledge Madeline Dawson for technical help and Katrina Granger, May Shen and Aishwarya Bhurke for data analysis and discussions. We are also grateful to Dr. Jason Bazil for Guinea Pig tissue and Dr. Linda Giudice for human tissue.

## DATA AVAILABILITY STATEMENT

Data generated in the manuscript is available upon reasonable request to the corresponding author.

---

**Supplementary Figure 1:**
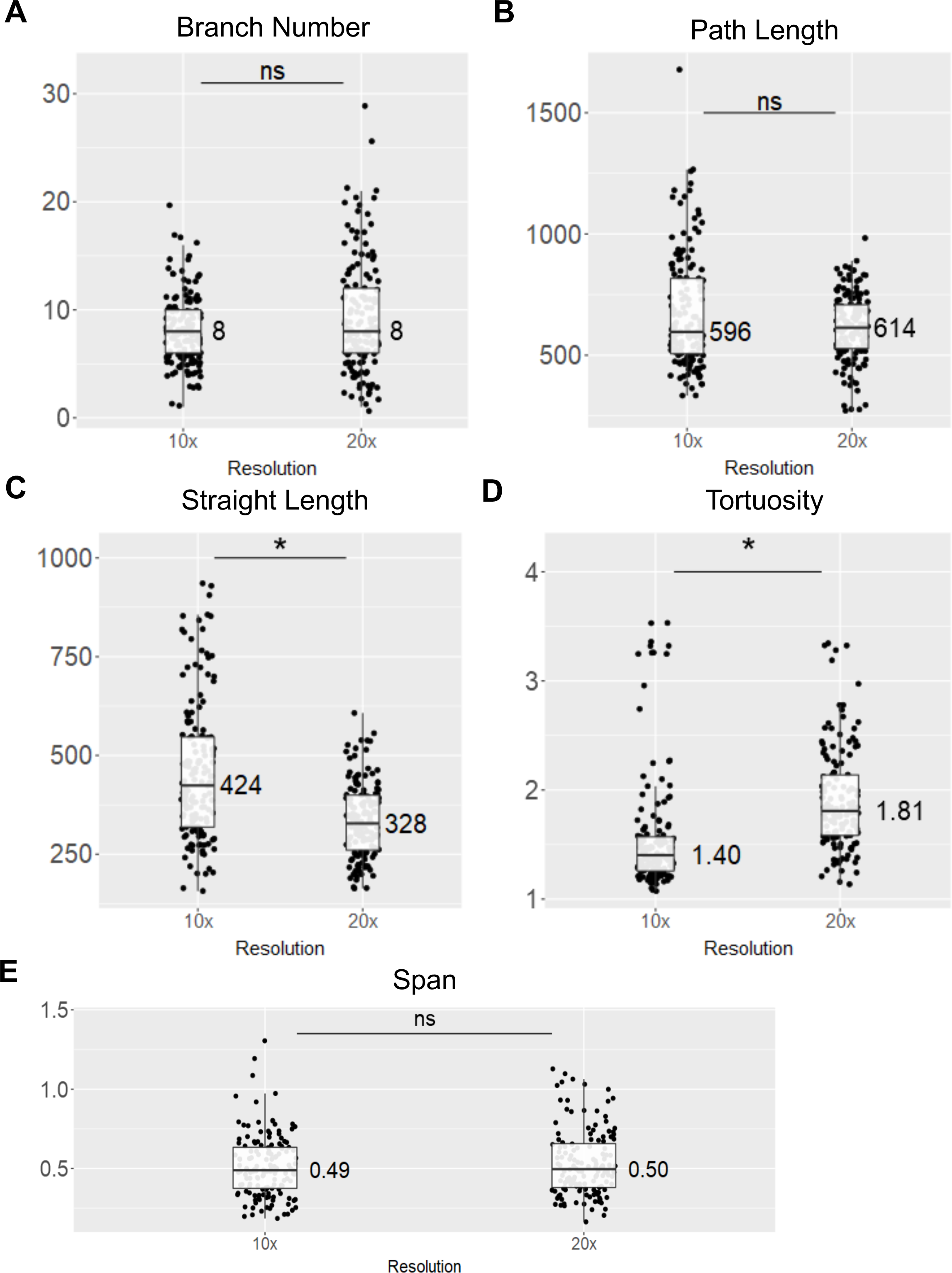
Metric Comparison Between Local Imaging in High Detail and Global Imaging in Low Detail. High-resolution imaging (20X) provides ground truth data and validates the utility of low-resolution imaging (10X). Metrics from high-resolution imaging are comparable to metrics from low-resolution imaging except straight length and tortuosity. (A) Comparison of branch number metric for glands at post-embryo-entry stage between 10x and 20x magnification. (B) Comparison of path length metric for glands at post-embryo-entry stage between 10x and 20x magnification. (C) Comparison of straight length metric for glands at post-embryo-entry stage between 10x and 20x magnification. (D) Comparison of tortuosity metric for glands at post-embryo-entry stage between 10x and 20x magnification. (E) Comparison of span metric for glands at post-embryo-entry stage between 10x and 20x magnification. * indicates p<0.05. ns = not significant; * = significant

**Supplementary Figure 2:**
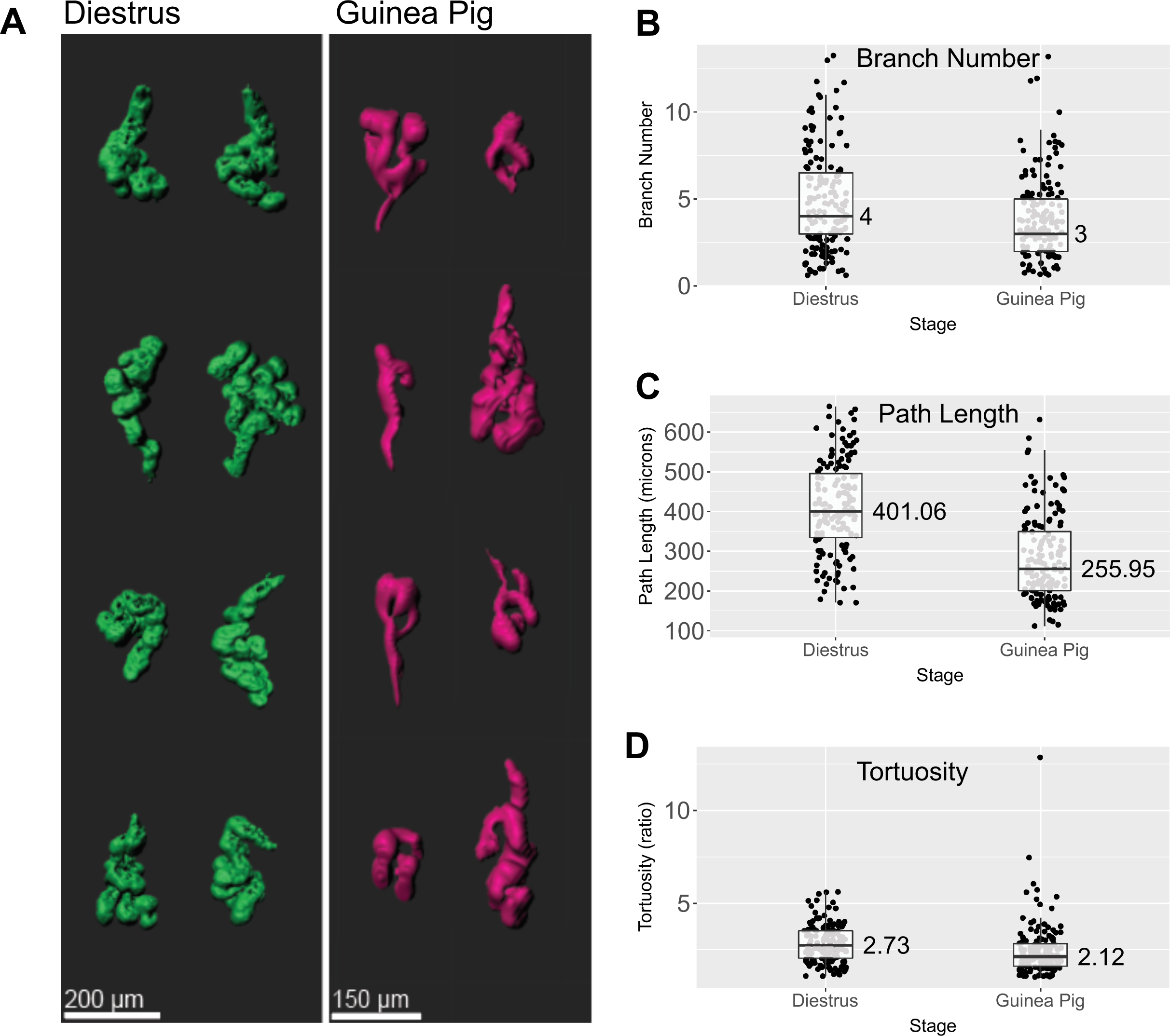
Metric Comparison Between Non-pregnant Guinea Pig and Mouse uterine glands. **(A)** 3D rendering of Gland surfaces from mice and guinea pigs. **(B)** Non-pregnant Guinea pig shows slightly decreased branching numbers compared non-pregnant mice. **(C)** Guinea pig glands are significantly shorter than mouse glands. **(D)** Tortuosity of guinea pig and mouse glands are relatively similar.

## Notes

### Competing Interest Statement

The authors have declared no competing interest.

