## Supplementary Methods for "Analysis Pipeline to Quantify Uterine Gland Structural Variations"

### **Additional Preprocessing Steps**

Preprocessing is required in some instances to counter the effects of poor antibody penetration and low signal to noise resulting in gland structure that is not well-defined (**Figure 1A**). As an initial step, Gaussian filters are applied in Imaris (see Image Processing: Smoothing: Gaussian Filter) to remove noise. Depending on the amount of noise, the sigma parameter can be increased, with the tradeoff that the gland structure becomes less defined. After Gaussian filtering, the image is exported from Imaris a TIFF file and is re-imported into MATLAB using the “imread_big” script (Ursell, 2022) in MATLAB. Optimally Oriented Flux (Law and Chung, 2008) or Vesselness3D (Jerman et al., 2016) filters can be applied to enhance the intensity of the gland and yield a well-defined structure. Optimally Oriented Flux functions best in low-noise contexts and preserves the branching structure of the gland better than Vesselness3D. Vesselness3D is employed in high-noise contexts where Gaussian filtering still leaves a large amount of noise. However, Vesselness3D causes gland branches that are close by to merge into single units.

All gland filtering steps are applied consistently across comparisons. The processed gland volumes are used as input to the Imaris Surfaces module to produce isosurface representations of the gland volumes. Distance transformation is applied to the gland surfaces (found under “Image Processing”, “Surfaces”, “Distance Transformation” once the surfaces are selected. The function will prompt the user on whether to calculate the metric as “Inside Surface” or “Outside Surface” select “Inside Surface”). This allows the user to encode voxels nearest to the center as highest intensity and voxels distant from the center as lower intensity. After all preprocessing, depending on the intended use case, the *Manual Filament, Automatic Filament, or Manual Trace* pipeline is executed (**Figure 1B**).

#### Manual Filament

The Imaris Filaments module is used to manually place points at the “root” of the gland where it attaches to the lumen and at each of the branches’ end points (**Figure 1B1, 1B2**). Imaris software computes the path between the gland “root” and the branch endpoint. The distance transformation ensures the calculated path is closest to the center of the gland lumen (**Figure 1C1**).

#### Automatic Filament

Gland surfaces are masked within Imaris (**Figure 1B3, 1B4**). All voxels outside the surface are encoded as zero intensity and all voxels inside the surface are encoded as 255 intensity, essentially binarizing the image (**Figure 1C2**). The volumes are exported from Imaris as a TIFF file and imported into MATLAB using the “imread_big” script (Ursell, 2022). After binarization, the gland volumes are skeletonized using the MATLAB Skeleton3D script (Kollmannsberger et al., 2017) to generate centerlines. This output is saved as a TIFF file using the “saveastiff” MATLAB script (Tak, 2022) and re-imported into the original Imaris file as an additional channel via the “Add Channels” option (found under “Edit” 🡪 “Add Channels”). Imaris Threshold Filaments are generated from the raw centerline image data output by the Skeleton3D script. Imaris Threshold Filaments are required in order to extract Imaris statistics and derive metrics from the raw data. Running Imaris Filaments module on the gland volumes directly is too cumbersome, uses too much graphic memory and often crashes the software. Skeletonization in MATLAB speeds up the process and makes the image analysis pipeline efficient.

#### Manual Trace

Manual Trace method requires intensive user input to trace the centerline through z-slices of the gland (**Figure 1B5, 1B6)**. This method is applicable for use cases where individual glands are difficult to separate from their neighboring glands when attempting to apply the Manual Filament method (**Figure 1C3**).

**Running the pipeline**

Users can download the entire codebase used to generate metrics from the Github space (<https://github.com/sameed-khan/filament-constructor>). To generate linear statistics and branch type classification, users can use the "manualFilamentsAllStats20x" MATLAB script and change the file paths to where their raw Imaris statistics CSV files are found on their local file system. To generate reorient and prominence statistics, users can use the "thresholdFilamentsAllStats" MATLAB script.

The supplementary code files can be used to generate metrics as follows:

Code 1 (path length, straight length, average branch length, tortuosity, span): **generateLengthStats**

Code 2 (branch type classification): **classifyBranchType**

Code 3 (prominence, reorient) - **generateReorientationThresholdFilaments**
