## Supplementary Code1 for "Analysis Pipeline to Quantify Uterine Gland Structural Variations"

```
function branchTypes = classifyBranchType(treeArray, maxPoints, outputPath)
% CLASSIFYBRANCHTYPE This function classifies the branch types for all branch
% points inside a Filament Surpass object (represented by treeArray)
%   @param treeArray: cell array; contains all FilamentTrees that you wish
%   to classify branches for
%   @OPTIONAL param outputPath: string; value defaults to false. If
%   provided, outputPath will generate a CSV file formatted as follows:
%       || Filament ID || Branch Point ID || Branch Type ||

    branchTypes = cell(maxPoints, 4);
    added_idx = 1;
    for i = 1:length(treeArray)
        tree = treeArray{i};
        % If Filament has less than 4 points, ID of last point is
        % considered to have no branches
        if length(tree.mapping) < 4
            branchTypes{added_idx, 1} = tree.filamentID;
            branchTypes{added_idx, 2} = tree.root.childrenByReference(1).id;
            branchTypes{added_idx, 3} = 'No Branches';
            branchTypes{added_idx, 4} = tree.root.childrenByReference(1).statistics('Pt Distance');
            added_idx = added_idx + 1;
            continue;
        end

        for j = 2:length(tree.mapping)
            fPoint = tree.mapping{j, 2};
            if fPoint.isTerminal()
                continue;
            end
            branchTypes{added_idx, 1} = tree.filamentID;
            branchTypes{added_idx, 2} = fPoint.id;
            branchTypes{added_idx, 4} = fPoint.statistics('Pt Distance');
            % Handle trifurcations
            if length(fPoint.childrenByReference) > 2
                bType = 'Trifurcation';
            elseif length(fPoint.childrenByReference) == 2
                childOne = fPoint.childrenByReference(1);
                childTwo = fPoint.childrenByReference(2);

                % Case 1: Two branch points
                if ~childOne.isTerminal() && ~childTwo.isTerminal()
                    bType = 'Structural Branch';
                % Case 2: Two terminal points
                elseif all([ childOne.isTerminal(), childTwo.isTerminal() ])
                    bType = 'Terminal Branch';
                % Case 3: One terminal point and one branch point
                else
                    bType = 'Side Branch';
                end
            else
                warning("(FilamentID: %s): Warning: Unhandled branch point case with only one child. Excluding...", num2str(tree.filamentID))
                continue
            end
            branchTypes{added_idx, 3} = bType;
```

```
            added_idx = added_idx + 1;
        end
    end
    branchTypes = branchTypes(1:added_idx-1, :);
    if nargin > 2
        tb = cell2table(branchTypes, 'VariableNames', {'filament_id',...
            'point_id','branch_class','pt_distance'});
        writetable(tb, outputPath);
    end
end
```
