## Supplementary Code2 for "Analysis Pipeline to Quantify Uterine Gland Structural Variations"

```matlab
function lenStats = generateLengthStats(treeArray, outputPath)
if nargin < 2
    outputPath = false;
end
lenStats = zeros(length(treeArray), 5);
for i = 1:length(treeArray)
    tree = treeArray{i};
    bP = [tree.root.statistics('Pt Position X'), tree.root.statistics('Pt Position Y'), tree.root.statistics('Pt Position Z')];
    tP = [tree.getExtendedTerminal().statistics('Pt Position X'), tree.getExtendedTerminal().statistics('Pt Position Y'), tree.getExtendedTerminal().statistics('Pt Position Z')];
    width = tree.statistics('Filament BoundingBoxOO Length B');
    lenStats(i,1) = tree.filamentID;
    lenStats(i,2) = tree.getExtendedTerminal().statistics('Pt Distance');
    lenStats(i,3) = norm(tP - bP);
    lenStats(i,4) = tree.getExtendedTerminal().statistics('Pt Distance') / norm(tP - bP);
    lenStats(i,5) = width / norm(tP - bP);
end
lenStats = array2table(lenStats, 'VariableNames', {'filament_id', 'path_length', 'straight_length', 'straightness', 'span'});
if outputPath
    writetable(output, sprintf('D:/Documents/Arora Lab Stuff/RAW_DATA/%s/%s/Output Statistics/Length_Stats_%i.csv', res,sample,i));
end
end
```
