## Supplementary Code3 for "Analysis Pipeline to Quantify Uterine Gland Structural Variations"

```
function reorient = generateReorientationThresholdFilaments(posPath, centerLine, orient, yBounds, outputPath)
addpath D:/Documents/MATLAB/utils
outputPathFlag = true;
if nargin < 5
    outputPathFlag = false;
end

% Read in uterine centerline approximation
utLine = readtable(centerLine, 'NumHeaderLines', 3, 'VariableNamingRule', 'preserve');
utLine = table2array(utLine(:, {'Position X', 'Position Y', 'Position Z'}));
utLine = sortrows(utLine,2);

termPoints = readtable(posPath, 'NumHeaderLines', 3, 'VariableNamingRule', 'preserve');
termPoints = termPoints(strcmp(termPoints.Type, 'Dendrite Terminal'),{'Pt Position X', 'Pt Position Y', 'Pt Position Z', 'FilamentID'});
fIDS = unique(termPoints{:, 'FilamentID'});

% Adjust y-axis values depending on location of cervix
if ~orient
    utLine(:,2) = (utLine(:,2) - yBounds(2)) * -1;
    termPoints{:,'Pt Position Y'} = (termPoints{:,'Pt Position Y'} - yBounds(2))*-1;
end
reorient = zeros(length(fIDS), 3);

% Calculate reorient by Filament
for i = 1:length(fIDS)
    %fprintf("Current ID: %i\n", fIDS(i));
    filPoints = termPoints(termPoints{:,'FilamentID'} == fIDS(i),{'Pt Position X', 'Pt Position Y', 'Pt Position Z'});
    pointArr = table2array(filPoints);

    % Find beginning pt by grabbing terminal pt closest to lumen line
    [~, dists, ~] = distance2curve(utLine, pointArr, 'spline');
    [~, min_idx] = min(dists);
    stPoint = pointArr(min_idx,:);

    % Find terminal point by grabbing pt with furthest distance
    [~, max_idx] = max(vecnorm(pointArr - stPoint, 2, 2));
    tPoint = pointArr(max_idx,:);

    % Compute statistic
    diff = tPoint(2) - stPoint(2);
    reorient(i, 1) = fIDS(i); % Filament ID
    reorient(i, 2) = diff; % reorientation statistic
    reorient(i, 3) = stPoint(2); % y-coordinate

end
reorient = array2table(reorient, 'VariableNames', {'filament_id', 'reorient', 'y_coord'});
if outputPathFlag
    writetable(reorient, outputPath);
end
end
```
